# Different effects of phytohormones on Fusarium head blight and Fusarium root rot resistance in *Brachypodium distachyon*

**DOI:** 10.1101/2020.06.26.173385

**Authors:** J F. Haidoulis, P. Nicholson

**Affiliations:** Department of Crop Genetics, John Innes Centre, Norwich Research Park, Norwich, NR4 7UH, England

**Keywords:** *Brachypodium distachyon*, Fusarium head blight, Fusarium root rot, Phytohormones

## Abstract

*Fusarium graminearum* is a devastating pathogen of small grain cereals causing both Fusarium head blight (FHB) and Fusarium root rot (FRR). Exogenous application of phytohormones has been shown to affect FHB resistance. In contrast to FHB, FRR remains poorly characterised and it is unknown whether phytohormones play similar roles in FHB and FRR. In this present study, *B. distachyon* floral tissues at mid-anthesis and root tissues from seedlings were exogenously treated with several phytohormones before inoculation with *F. graminearum*. The canonical defence-associated phytohormones had differing effects on FHB and FRR. Salicylic acid (SA) significantly increased susceptibility to FRR but not to FHB while jasmonic acid (JA) and ethylene increased resistance to FRR but increased susceptibility to FHB. Additionally, the growth-associated phytohormones auxin and cytokinin significantly increased resistance and susceptibility, respectively, to both diseases. This study is the first to compare phytohormone effects between FHB and FRR in the same host.

**Highlight:** The tissue-dependent effects of defence phytohormones and tissue-independent effects of development phytohormones on *F. graminearum*-induced Fusarium head blight and Fusarium root rot diseases in the model cereal *Brachypodium distachyon*.

## Introduction

*Fusarium graminearum* is one of the most important plant pathogens in the world (Dean et al., 2012), affecting small grain cereals such as bread wheat (*T. aestivum*), barley (*H. vulgare*), and rye (*S. cereale*). *F. graminearum* can infect almost the entire plant (Miedaner, 1997) causing a range of diseases. The most well-known and economically important disease is Fusarium head blight (FHB). Other diseases include Fusarium crown rot, Fusarium root rot (FRR), and seedling blight. Plant pathogens often adopt specific trophic lifestyles. Biotrophs obtain nutrients from living tissue, necrotrophs from dead tissue, and hemibiotrophic pathogens adopt an early biotrophic phase that is followed by a necrotrophic phase sometime later in infection (Glazebrook, 2005, Zeilinger et al., 2016). *F. graminearum* is generally considered a facultative hemibiotroph in the developing cereal spike (Jansen et al., 2005, Boddu et al., 2006). Infection patterns of floral tissues show that *F. graminearum* grows as a biotroph at the advancing hyphal front with necrotrophic feeding behind this, where hyphae are ramifying through tissues already killed by the fungus (Brown et al., 2010). Current FHB control methods rely on the effective use of fungicides and genetics resistance. However novel control mechanisms are required.

Fungal root diseases are widespread in wheat and barley fields and are becoming more prevalent with the increased use of cereal crop rotation and no tillage practices (Cook, 2001). FRR is caused by several Fusarium species including *F. graminearum*. Colonisation and sporulation can occur rapidly (Wang et al., 2015) causing root browning and necrosis (Beccari et al., 2011). FHB and Fusarium crown rot can also manifest from FRR infection due to systemic migration via the vascular system (Beccari et al., 2011, Wang et al., 2015). Ultimately this results in reduced root, shoot length, biomass and yield loss (Mergoum et al., 1998, Beccari et al., 2011, Wang et al., 2015). Control of FRR is also difficult due to limited effectiveness of fungicides, lack of genetic resistance, and persistence in the soil for many years facilitated by the predominantly saprotrophic lifestyle of *F. graminearum* (Cook, 2001). Furthermore, FRR is not as well characterised as FHB due to the difficulty of research on root diseases.

*Brachypodium distachyon* (purple false brome) is a valuable model for small-grained cereals as it is a small inbreeding annual with short generation times, minimal growth requirements, and abundant genetic resources (Draper et al., 2001, Vogel et al., 2006, Vogel et al., 2010, Brkljacic et al., 2011, Kellogg, 2015, Scholthof et al., 2018). *B. distachyon* shoots and roots show high anatomical and developmental similarities to wheat (Draper et al., 2001, Watt et al., 2009, Opanowicz et al., 2011, Chochois et al., 2012), and hormone signalling pathways appear conserved (Goddard et al., 2014, Powell et al., 2017). Importantly, *B. distachyon* is an excellent model for investigating FHB and FRR as both roots and florets are susceptible to *F. graminearum* infection (Peraldi et al., 2011, Pasquet et al., 2014).

Phytohormones are critical components of plant defence signalling and their impact on *F. graminearum* resistance response has been investigated in numerous studies on *A. thaliana*, wheat, and barley. Exogenous application of salicylic acid (SA) or methyl salicylate has been shown to have positive effects on FHB resistance in wheat (Makandar et al., 2011, Qi et al., 2012, Sorahinobar et al., 2016) with direct effects on *in-vitro* growth (Qi et al., 2012). Application of methyl salicylate also showed positive effects on wheat leaf resistance to *F. graminearum* (Ameye et al., 2015). Jasmonic acid (JA) and methyl-jasmonic acid application was also reported to have a positive effect on FHB resistance in wheat (Li and Yen, 2008, Qi et al., 2016, Sun et al., 2016). In contrast Makandar and colleagues (2010) found that methyl-jasmonic acid compromised resistance during early infection but had positive effects during late infection in *A. thaliana* (Makandar et al., 2010). Similar results were reported in wheat where pre-infection application of methyl-jasmonic acid promoted susceptibility, whereas post-infection application promoted resistance of wheat leaves to *F. graminearum* (Ameye et al., 2015). JA was also found to reduce *F. graminearum* growth in vitro (Qi et al., 2016). The precise role of ethylene on resistance to Fusarium remains unclear. Compounds that produce ethylene have been reported to have positive effects (Li and Yen, 2008, Foroud et al., 2018), negative effects (Chen et al., 2009), and no significant effect (Sun et al., 2016) on Fusarium resistance.

More recently other important hormones were shown to have roles in defence and contribute significantly to the infection response. Exogenous application of gibberellic acid reduced FHB spread in wheat heads (Buhrow et al., 2016), whereas exogenous application of abscisic acid negatively affected resistance to FHB (Buhrow et al., 2016, Qi et al., 2016). Exogenous application of brassinosteroid had positive effects on resistance (Ali et al., 2013) while brassinosteroid receptor mutants in which brassinosteroid levels are believed to be increased, exhibited enhanced resistance to *Fusarium culmorum* (Goddard et al., 2014). Finally, the auxin indole-3-acetic acid (IAA) had positive effects on barley FHB (Petti et al., 2012), and was even found to reduce Fusarium growth *in vitro* (Luo et al., 2016). There are hormones that have not been investigated regarding exogenous application and response to *F. graminearum* infection. These include cytokinins which have been implicated in plant defence (Choi et al., 2010, Choi et al., 2011, Albrecht and Argueso, 2017) and the recently classed non-protein amino acid and signalling molecule 3-aminobutryric acid (BABA) (Cohen, 2001, Jakab et al., 2001, Cohen, 2002, Thevenet et al., 2017).

The reported effects of phytohormones on resistance to *F. graminearum* is complex or in some cases contradictory. Most studies have focused on FHB with a lack of research on *F. graminearum*-induced FRR pathogenesis. The main aim of this research was to investigate the effect of several phytohormones on resistance to FRR and compare these with the effects of the same compounds on FHB. We then generated a model for how each phytohormone influences resistance or susceptibility to the two diseases in *B. distachyon*.

## Materials and Methods

### Plant Material and Growth Conditions

The *Brachypodium distachyon* line Bd3-1 was obtained from the John Innes Centre, Norwich, UK. To soften the floret, seeds were soaked in water for 10-30 min. Subsequently, the lemma and palea were peeled off the individual seeds. Seeds were placed between two layers of dampened filter paper (9 cm, Sartorius Grade 292) with 5 ml sterile water, and were stratified at 5 °C for five days in the dark, and incubated for one day at 22 °C (16h/8h light/dark photoperiod, variable humidity) in controlled environment growth cabinets (Peraldi et al., 2011). For FHB assays, peeled Bd3-1 seeds were sown in 50% Peat/Sand and 50% John Innes mix 2 (two seeds per pot). Plants were kept for four weeks at 22°C (20h/4h light/dark photoperiod, 70% humidity) in controlled environment growth cabinets or rooms (Peraldi et al., 2011). The *B. distachyon* line Bd21 was obtained from BASF SE Agricultural Centre Limburgerhof, Germany, For the FHB assay, approximately five Bd21 unpeeled seeds were grown for 6 weeks at 20h/4h (day/night) in a glasshouse (Autumn conditions).

### Preparation and Maintenance of *Fusarium graminearum*

The *Fusarium graminearum* isolate PH1 was used for all experiments. In order to produce mycelial inoculum for FRR assays, *F. graminearum* was maintained on approximately 20 ml potato dextrose agar (PDA) in 9 cm diameter plastic Petri-dishes in a controlled growth cabinet at 22°C under 16h/8h light/dark photoperiod. Conidial *F. graminearum* inoculum was produced in Mung Bean broth as reported previously (Makandar et al., 2006). After one week, conidia were harvested, washed and concentration adjusted to 1 × 10^6^ conidia/ml. Inoculum was stored at 5 °C for up to two weeks before inoculation. For Bd21 FHB assay, *F. graminearum* isolate Li600 was grown alternating on malt agar and oatmeal agar for two weeks and conidia were harvested in 10 ml of water amended with 0.05 % Tween 20 and concentration adjusted as above.

### Fusarium Root Rot Assays with Chemical Amendment

Aspects of the FRR assay were derived from (Peraldi et al., 2011, Goddard et al., 2014) and modified for chemical amendment experiments. A sterile 9 cm^2^ filter paper square (cut from Chromatography Paper 46 cm x 57 cm from Slaughter Ltd R & L) was placed on square plastic square Petri-dishes (10 cm^2^) containing 50 ml autoclaved 0.8 % agar (Fischer Science). Under sterile conditions, ten cold-stratified and germinated Bd3-1 seeds were placed on the filter paper. A minimum of 30 seedlings was used for each treatment. All plates were placed, angled at 70° from the horizontal to ensure uniform downward root growth, in covered plant propagators containing wetted paper towel to maintain high humidity (Supp Mat. Fig. x). Plant propagators with square Petri-dishes were incubated at 22 °C (16h/8h light/dark photoperiod, variable humidity) in controlled environment growth cabinets. After three days, the filter paper with seedlings attached was carefully transferred to different square Petri-dishes containing 0.8% agar amended with phytohormone or control solvent alone and returned to controlled environment growth cabinets. All compounds were ordered from Merck/Sigma-Aldrich UK unless otherwise stated. The final concentration for SA, JA, ACC, and 3-aminobutanoic acid (BABA) were primarily derived from dilution series experiments in combination with information from (Kakei et al., 2015) for the SA concentration, (Dr. Antoine Peraldi, unpublished) for the JA concentration, (Peraldi, 2012, Van de Poel and Van Der Straeten, 2014) for the ACC concentration, and (Cohen, 2002, De Vleesschauwer et al., 2010) for the BABA concentration. The concentration for IAA was derived from (Kakei et al., 2015) whereas trans-Zeatin was derived from (Großkinsky et al., 2013, Kakei et al., 2015). The concentrations for the 1-napthaleneacetic acid (NAA) and kinetin were derived from the concentrations of IAA and trans-Zeatin respectively. The compounds ACC, IAA – sodium salt, kinetin, and BABA were dissolved in water. The compounds SA, JA and NAA were dissolved in ethanol and trans-zeatin was dissolved in DMSO. All non-water solvent concentrations in final treatment were at or below 0.1 %. The same concentration of solvent was applied to respective control treatment 0.8 % agar plates.

Roots were inoculated six hours after transfer to hormone amended medium. Inoculum was prepared by blending mycelium and PDA from one-week old cultures (adding 1 ml water per PDA plate). Subsequently, approximately 0.1 ml to 0.2 ml of homogenized mycelial slurry was transferred with a 10 ml syringe onto the root tip. Inoculated plants were then incubated at 22 °C (16h/8h light/dark photoperiod, variable humidity) in controlled growth cabinets. The slurry was removed when necrosis was visible at the root tip, often at 1-day post inoculation (dpi). Roots were photographed at intervals to monitor disease development. For repeated independent experiments with different measurement dates, the days were combined and denoted as ‘Score Date’ 1 to 3 with each measurement at 3-day intervals (Fig. 1B, Fig. 1C).

**Fig. 1:**
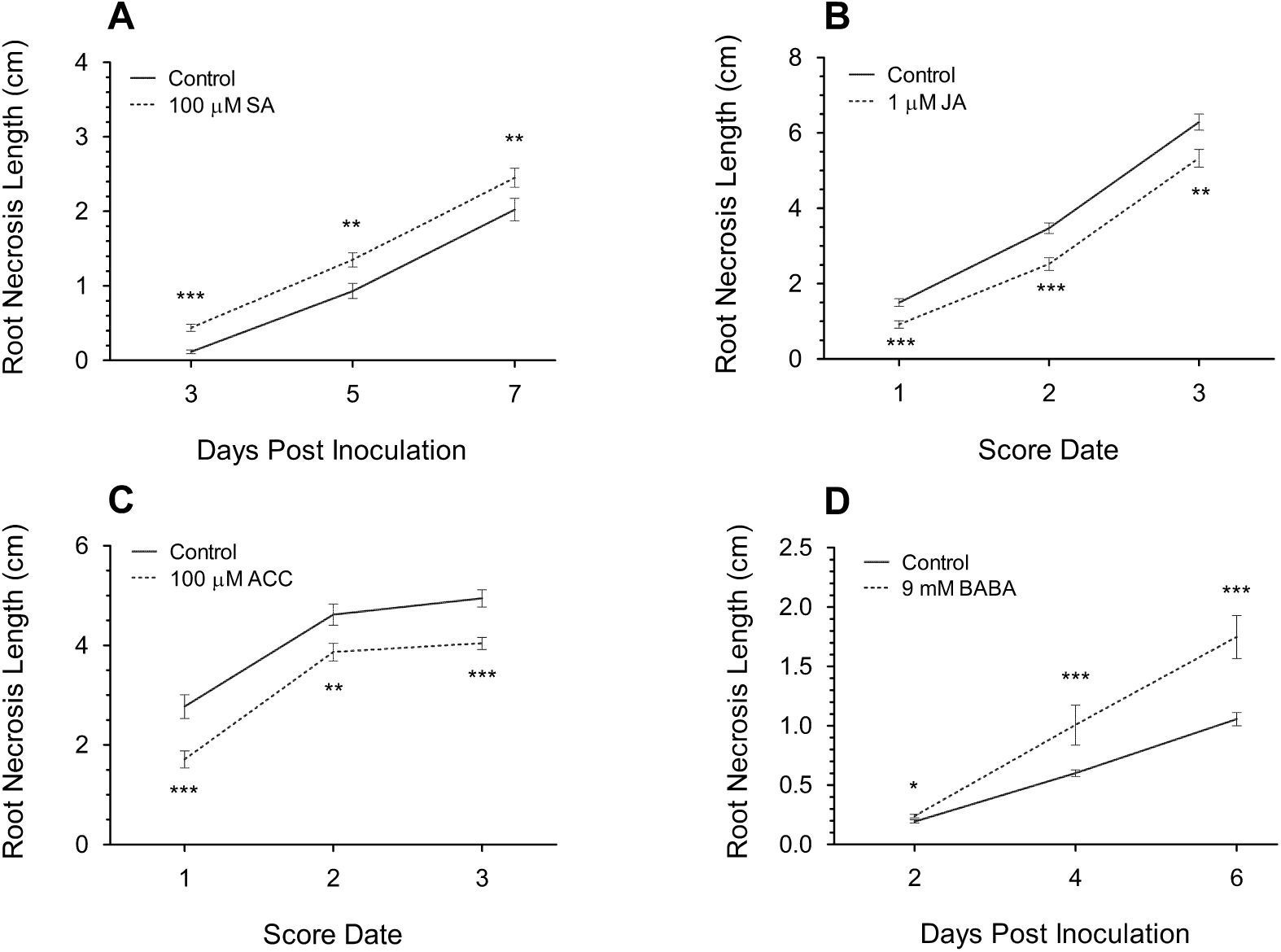
The change in *F. graminearum*-induced FRR necrosis after application of defence-related hormones 100 µM SA (A), 1 µM JA (B), 100 µM ACC (C), and 9 mM BABA (D) on Bd3-1 seedling roots. Each data point is the mean root necrosis length ± SE from two (A, B, C) or three (D) independent experiments, except score date 3 from C which was from one independent experiment. Score Date numbers (3-day intervals between each number) for B and C are the combined dpi from different experiments. Significance levels: ** p < .01, *** p < .001 compared to control.

### Fusarium Head Blight Assays with Chemical Pre-Treatment

Numerous features of the FHB assay were derived from (Peraldi et al., 2011). With Bd3-1 FHB assays, once extruding anthers were visible around mid-anthesis, the entire Bd3-1 plant was sprayed with 50 ml of phytohormone or solvent with 0.05% Tween 20 onto a tray of 10 to 11 pots. All compounds were ordered from Merck/Sigma-Aldrich UK unless otherwise stated. The concentrations used for BABA and trans-Zeatin were derived from those used in FRR assays, the SA concentrations was obtained from previous publications (Mandal et al., 2009, Makandar et al., 2011, Sorahinobar et al., 2016, Makandar et al., 2010) and JA and ACC were experimentally determined. ACC and BABA were dissolved in water, SA and JA in ethanol, and trans-Zeatin in DMSO. The final concentration of non-water solvents was kept at or below 0.2 %. The same concentration of solvent was applied to respective control treatment groups. Twenty-four hours later the soil was watered, and Bd3-1 spikes were sprayed to run-off with conidia of *F. graminearum* (0.25-1 × 10^6^) amended with wetting agent (0.05 % Tween 20). A control pot without inoculum was included for each treatment. Approximately 30 ml of inoculum was sprayed onto a total of 20 plants’ spikelets per treatment. Spraying was performed immediately before the dark period and pots were then collectively held in a large plastic humidity chamber to maximise humidity for three days 22°C (20h/4h light/dark photoperiod). Symptoms were scored every four days by counting the number of infected florets per spike. For repeated independent experiments with different measurement dates, the days were combined to ‘Score Date’ 1-3 with each measurement at 4-day intervals (Fig. 3B, Fig. 3D).

**Fig. 2:**
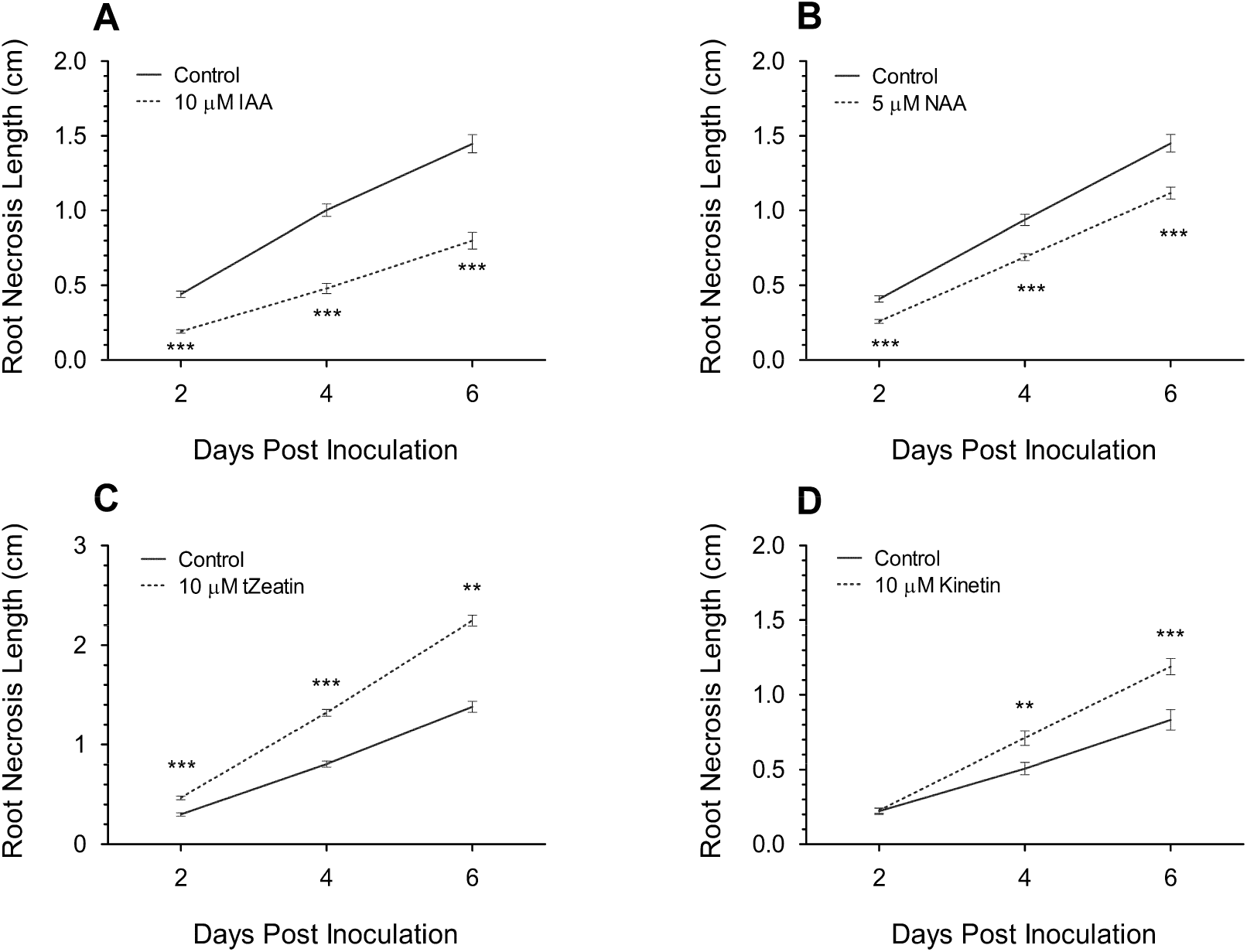
The change in *F. graminearum*-induced FRR necrosis after application of development-related auxins 10 µM IAA (A) and 5 µM NAA (B), and the cytokinins 10 µM trans-Zeatin (C) and 10 µM kinetin (D) on Bd3-1 seedling roots. Each data point is the mean root necrosis length ± SE from three (C), two (A and B) or one (D) independent experiments. Significance level ** p < .01, *** p < .001 compared to control. Visual difference for A in Supp. Fig. 4 and C in Supp. Fig. 5.

**Fig. 3:**
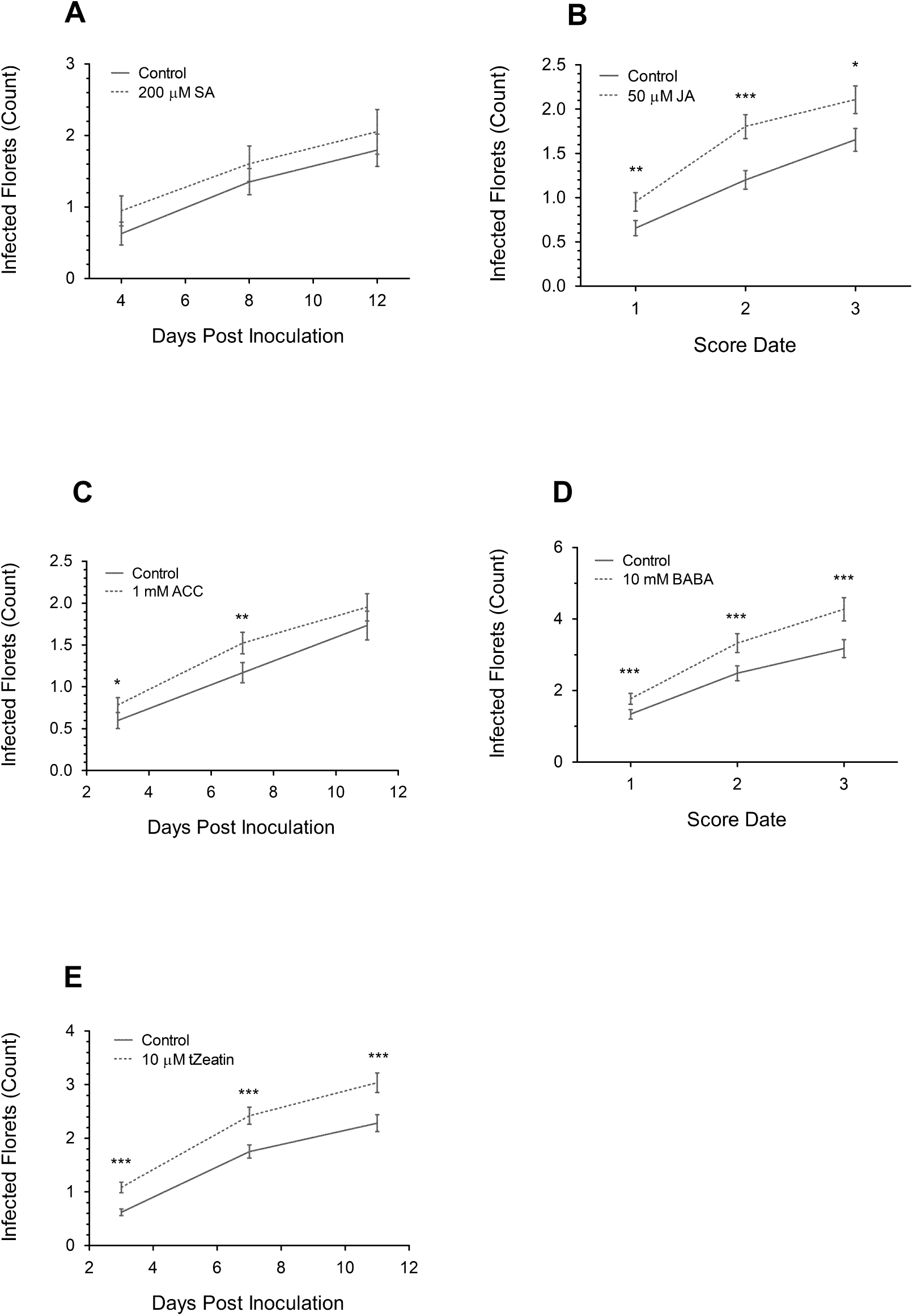
The change in number of *F. graminearum*-infected florets after pre-application of 200 µM SA (A), 50 µM JA (B), 1 mM ACC (C), 10 mM BABA, (D) and 10 µM trans-Zeatin (E) on Bd3-1. Each data point is the mean number of florets infected ± SE from one (A), two (B, C, E) or three (D) independent experiments. Score Date numbers (4-day intervals between each number) for B and D are the combined dpi from different experiments. Significance levels: * p < .05, ** p < .01, *** p < .001 compared to control.

Some experiments were performed at BASF SE using Bd21 instead of Bd3-1. Bd21 and Bd3-1 are both highly susceptible to FHB (Peraldi et al., 2011). For the Bd21 auxin trial, the entire plant was sprayed with respective compounds until run-off. The auxin concentration was experimentally determined (Unpublished data). The final solvent concentration for each auxin treatment with Bd21 including the control treatment was 5 % acetone. After 24h, a working concentration of 5×10^5^ *F. graminearum* LI600 was prepared in 0.05 % tween 20 water, and 175 ml of inoculum was evenly sprayed above all plants. The base matting was watered until run-off and plants were encased in plastic covers for six days in an elevated humidity glasshouse room (Autumn conditions) before scoring.

### Statistics

All statistical tests were performed using the software package GENSTAT v.19.1.0.21390 (VSN international Ltd). A Generalised liner model with an accumulated analysis of variance (ANOVA) was used for all experiments using a normal distribution for FRR assays or log-linear modelling for FHB assay count data. Individual time points were analysed separately. All data was untransformed except for a square root transformation for Fig. 1A at 7 dpi, Fig. 1C at score date 1, Fig. 2A at 2 dpi, Fig. 2C at 4 dpi, Fig. 2D at 2dpi and 4dpi. A log10 transformation was used for Fig. 2C at 2dpi. A few ANOVA’s for combined independent experiments showed a significant interaction between the independent experiment and the hormone treatment at specific time points: Fig. 1A at 3 dpi, Fig. 1B at Score Date 3, Fig. 1D at 2 dpi, Fig. 2A at 2 dpi, Fig. 2B at 2 dpi, Fig. 3C at 11 dpi, and Fig. 3E at 7 dpi and 11dpi (p < .05). However for all the remaining time points, no such interaction was observed (p > .05). For Bd21, a randomised complete block design was used from the R package ‘agricolae’ and plant pots were randomized before chemical application. Microsoft Office was used for writing, data collection, diagrams, and analysis. All graphs were prepared using Graphpad Prism (V5.04).

## Results

### The Differential Effect of Plant Hormones on Fusarium Root Rot Disease Progression

The SA amendment significantly increased root necrosis length (RNL) at all time points (Fig. 1A), while amendment with JA (Fig. 1B), or the ethylene precursor (ACC) (Fig. 1C), significantly reduced RNL at all time points (p < .01). Interestingly, the difference in RNL over time for all three compounds remained similar with the regression lines remaining parallel rather than diverging over time. The compound BABA had the greatest effect in increasing RNL compared to other defence-related hormones (Fig. 1D). The differential RNL symptoms resulting from pre-application of BABA increased over time with application leading to near doubling of necrosis at 4 dpi and 6 dpi relative to the control (p < .001). Overall exogenous application of defence-related hormones had a substantial effect on FRR disease severity but whether the effect on resistance was positive or negative was dependant on the phytohormone.

Exogenous application of IAA resulted in the most pronounced decrease in RNL compared to the control at all time points (p < .001) since FRR symptoms were reduced to approximately half those in control treatments (Fig 2A). Like IAA, the synthetic auxin NAA also reduced RNL at all time points (p < .001, Fig 2B) but auxin induced FRR resistance was most pronounced with IAA. With both auxins, the increased resistance was observed as early as 2 dpi yet the regression lines diverged over time with a greater reduction in RNL at later time points (Supp. Fig. S1). Overall, auxin treatment produced a very positive effect on resistance towards FRR.

The cytokinin trans-Zeatin increased RNL at all time points (p < .001, Fig. 2C) and the differential increased over time. The extent of necrosis was twice that in the control by 6 dpi (Supp. Fig. S2) leading to the most severe symptoms observed with any of the phytohormone treatments. The cytokinin kinetin exhibited no significant effect on symptoms at 2dpi (p = .785) but increased the rate of RNL so that the differential increased over time ((4 dpi and 6 dpi p < .01, p < .001 respectively) Fig. 2D). Strikingly the effect of cytokinin amendment increased dramatically over time, with the effect being even greater than that observed for the positive effect of auxins. Overall cytokinin amendment appeared to have a very negative effect on FRR resistance which became more pronounced as the disease progressed.

The phytohormones abscisic acid, gibberellic acid, and the brassinosteroid epibrassinolide were also tested on FRR (Supp. Fig. S3A and S3B and Supp. Fig. S4). Gibberellic acid and epibrassinolide had no significant effect on FRR symptoms (Supp. Fig. S3A and S3B) while abscisic acid induced extensive root discolouration that was indistinguishable from FRR symptoms and so prevented assessment (Supp. Fig. S4).

### Most Plant Hormones Promoted Susceptibility to Fusarium Head Blight

In order to compare the effects of hormones on FRR to those on FHB, the same hormones were applied to the entire *B. distachyon* plant and subsequently the spikes were inoculated with *F. graminearum* conidia. Unless otherwise stated, the effects described are from a single dose of phytohormone pre-treatment as opposed to continual exposure as during the FRR assays. As a result, often a higher concentration of phytohormone was applied to maximise the dose of phytohormone received by the plant.

Although pre-inoculation treatment with SA marginally increased FHB symptoms (Fig. 3A), the increase was not statically significant at any time point (p > .05). In additional experiments, SA was applied repeatedly (four applications) before, during, and after inoculation on Bd3-1 but again, no significant effect on resistance was observed (Supp. Fig. S5). Pre-inoculation treatment with JA resulted in an increase in the number of infected florets at all time points (p < .05, Fig. 3B). Similarly, pre-treatment of ACC (Fig. 3C) also significantly increased the number of infected florets at all time points (p < .05), but the effect on symptoms diminished over time and by 11 dpi, the effect was not significant (p > .05). Pre-inoculation application of BABA significantly increased the number of infected florets at all time points (p < .001) (Fig. 3D) and application of cytokinin trans-Zeatin caused the largest increase in infected florets relative to the control treatment of any of the compounds examined (Fig. 3E) (p < .001). With both BABA and trans-Zeatin, the differential in the number of infected florets relative to the control treatments increased after the first score date although the divergence from the controls was not as great as observed in the FRR assays where the phytohormone amendment was present throughout the course of the experiment. Overall pre-inoculation application of all the phytohormones except for SA promoted FHB susceptibility leading to a significant increase in the number of florets exhibiting symptoms.

Using Bd21 instead of Bd3-1, two concentrations of IAA and NAA were exogenously applied to Bd21 plants before inoculation with the isolate *F. graminearum* LI6000 (Fig. 4). All auxin pre-inoculation applications resulted in a reduction in the number of infected florets. Pre-inoculation application of synthetic auxin NAA at 8 mM resulted in the most significant decrease in the number of infected florets (p < .001) and even 4 mM NAA significantly decreased the number of infected florets (p < .01). IAA appeared to be less effective than NAA since 8 mM IAA showed a similar reduction in infected florets as 4 mM NAA (p < .01) and at 4 mM IAA the reduction in the number of infected florets was not significant (p = .33). Overall, these data indicate that auxins have a positive effect on FHB resistance when applied at relatively high concentrations prior to inoculation.

**Fig. 4:**
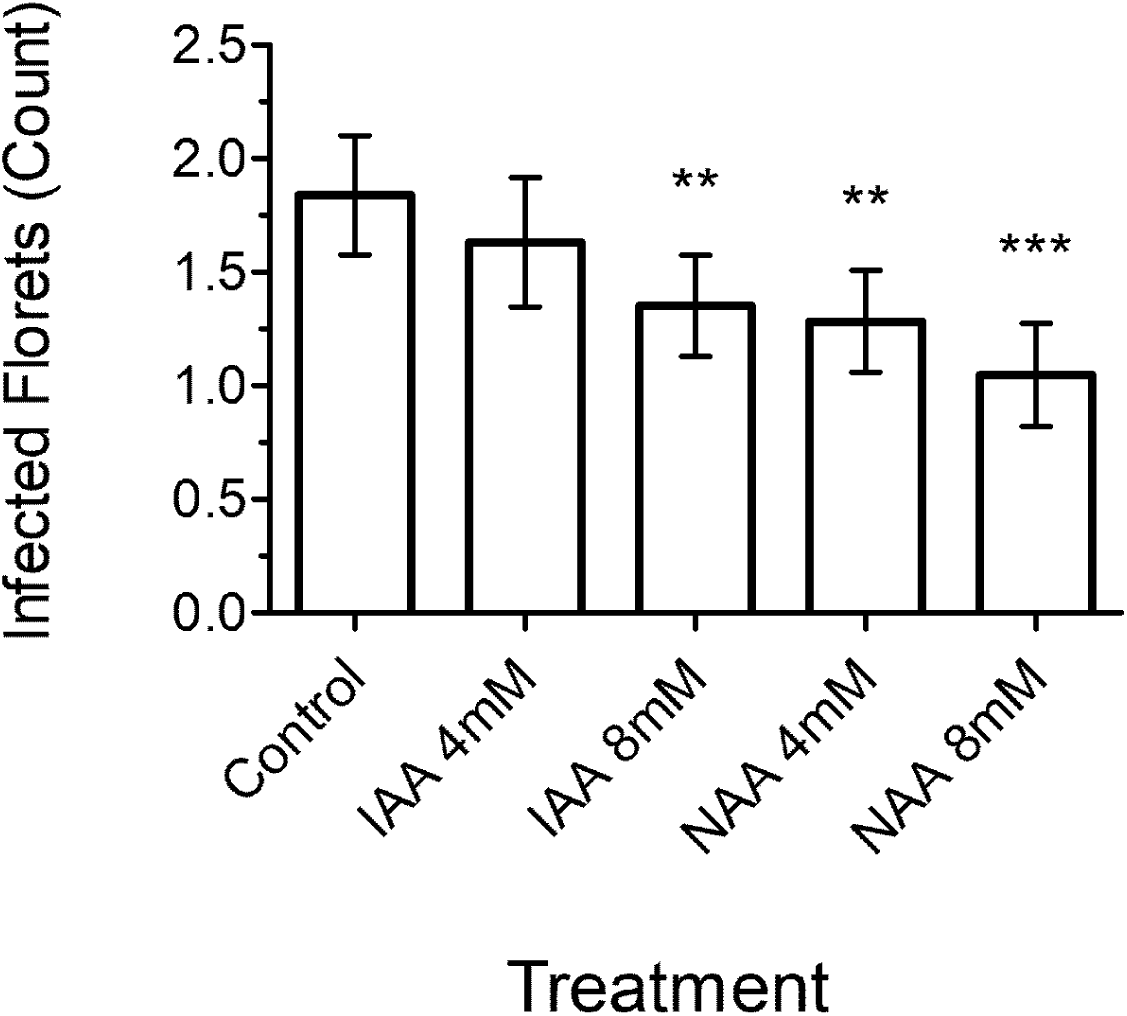
The number of *F. graminearum*-infected florets at 8 dpi after pre-application of IAA or NAA on Bd21. Each bar is the mean number of florets infected ± SE from one experimental replicate. Significance levels: ** p < .01, *** p < .001 compared to control.

To ascertain whether the effects of phytohormones on FRR reflected an altered plant response or an effect on the fungus itself, hormones at the same concentrations as used in the experiments were applied to *F. graminearum* growth medium to measure any changes in growth over time. Compared to the respective control treatments there was no significant effect on *F. graminearum* growth of any of the tested phytohormone (Supp. Fig. S6). Due to the high concentrations of auxins tested for FHB, a different assay was performed. Auxins or solvent controls were applied to filter disks at four equidistant points from a mycelial plug of *F. graminearum* PH1 growing on PDA. There was no difference in mycelial growth near any of the auxin-treated filter disks (Supp. Fig. S7).

## Discussion

Given some of the proposed differences between *F. graminearum* infection of floral and root tissues, the effect of a range of phytohormones on FRR was assessed and compared to their effect on FHB. Research into the role of exogenous phytohormones on FHB resistance has been investigated in several plant species. SA was found to increase resistance to FHB in wheat and Arabidopsis (Makandar et al., 2011, Qi et al., 2012, Sorahinobar et al., 2016). We were not able to replicate the positive FHB effect of SA (Fig. 3A) despite using similar concentrations to those used previously. We speculate that the different results are due to alternative SA application methods. In earlier studies, SA (200 µM) was applied as a soil drench (Makandar et al., 2011, Sorahinobar et al., 2016). This application method may have more pronounced effects on resistance due to initial uptake in the roots as opposed to florets and because the phytohormone remained in contact with the roots throughout the experiment. Furthermore prolonged exposure to SA may influence JA/ethylene dependent Induced systemic resistance (ISR) (Pieterse et al., 2014) which might impact the resistance response to FHB.

Our results on the effect of JA application on FHB (Fig. 3B) support some studies (Makandar et al., 2010), but contradict others (Li and Yen, 2008, Qi et al., 2016, Sun et al., 2016). The reasons for these differences are unclear. They may reflect the use of a higher concentration of JA in the present study than was used in some previous reports (Sun et al., 2016, Qi et al., 2016) or they may be due to differences in the timing of treatment (Makandar et al., 2010). The difference may also be due to the application method. Qi and colleagues (2016) point-inoculated individual florets with *F. graminearum* rather than spraying the head (Qi et al., 2016). Injecting high levels of inoculum into florets may have resulted in a by-passing or severe shortening of the biotrophic phase of infection/colonisation (Jansen et al., 2005, Brown et al., 2010). Similar to JA, our results with ACC (Fig. 3C) support the findings from some studies (Chen et al., 2009), but not those of others (Li and Yen, 2008, Sun et al., 2016, Foroud et al., 2018). Ethylene can affect host resistance via synergism with classic defence hormones (Pieterse et al., 2012) or via accelerated senescence (Abeles et al., 2012, Häffner et al., 2015). As a result, the outcome is very sensitive to several factors such as timing of exogenous application, tissue type, and the type of invading pathogen (van Loon et al., 2006). Differences in the experimental procedures may also affect important factors for ethylene response. In some reports detached head assays were performed (Chen et al., 2009, Foroud et al., 2018) which might influence senescence processes as well as defence responses. Ethephon (releases ethylene gas) was used instead of ACC as the ethylene treatment in some studies (Li and Yen, 2008, Chen et al., 2009, Sun et al., 2016). Although Foroud and colleagues (2018) found that ethephon and ACC had similar effects on FHB in most wheat varieties tested (Foroud et al., 2018). The role of ethylene is further complicated given evidence that some isolates of *F. graminearum* and *F. culmorum* can produce ethylene *in vitro* (Haidoulis et al. unpublished data), however it is uncertain whether ethylene is produced *in planta*.

We found that auxin, (both IAA and NAA) greatly improved FHB resistance (Fig. 4). This results supports evidence by Petti and colleagues (2012) who found that exogenous IAA application reduced yield loss and general symptoms of *F. culmorum*-induced FHB in barley (Petti et al., 2012). However, the mechanism for the effects of auxin in FHB response are not clear given that *F. graminearum* can also produce IAA (Luo et al., 2016).

Overall, our results using *B. distachyon* as a model for investigating FHB responses were broadly in line with those from studies on wheat, barley and Arabidopsis. Similar trends were observed in the response to auxins, JA, and ethylene as reported in FHB of wheat and barley. Wheat and *B. distachyon* have been show to share a common defence transcriptome response following Fusarium crown rot infection caused by *F. pseudograminearum* (Powell et al., 2017). Furthermore *B. distachyon* and barley were found to share common brassinosteroid-associated defence responses through the use of brassinosteroid receptor (*bri1*) mutants in the two species (Goddard et al., 2014). This suggests the observed effects of phytohormones on FHB in *B. distachyon* can be effectively translated to wheat FHB responses.

The pathology of *F. graminearum* in root tissues is poorly characterised, but evidence suggests that the fungus rapidly penetrates roots and exhibits specific stages of infection (Wang et al., 2015). Investigation of exogenous phytohormones on FRR response revealed that FRR was much more responsive than FHB in the response to infection. This is likely due to the continual exposure during FRR assays as opposed to the single application used for FHB assays. An example of this was observed JA (Fig. 3B) and ACC (Fig. 3C) that displayed the most pronounced effect on FHB susceptibility at earlier time points suggesting a short-lived response.

The classic defence hormones SA, JA, and ethylene (ACC) exhibited different effects on FRR and FHB (Fig. 5). Despite the non-significant FHB response to SA, SA increased FRR susceptibility as early as 1 dpi (Fig. 1A). This difference may reflect the differences in the levels of phytohormones in different tissues. Endogenous free SA levels were found to be substantially higher in rice (*O. sativa*) floral tissues compared to root tissues (Chen et al., 1997). If this is also true for *B. distachyon*, exogenous SA application might be expected to have more impact on the endogenous SA profile in roots than florets resulting in a more pronounced change in resistance in FRR compared to FHB. The similar response between JA and ethylene in FHB and FRR supports the evidence that JA and ethylene often function similarly in defence (Pieterse et al., 2012). However, both JA and ethylene had contrasting effects on FRR and FHB, increasing resistance to the former and susceptibility to the latter (Fig. 1B-C and Fig 4B-C). The SA pathway is generally considered to promote resistance to biotrophic pathogens and susceptibility to necrotrophic pathogens whereas JA and ethylene pathways generally promote resistance to necrotrophic pathogens and susceptibility to biotrophic pathogens (Glazebrook, 2005, Bari and Jones, 2009, Pieterse et al., 2012). In both FHB and FRR, hormones were applied before inoculation and would most likely influence events relating to the biotrophic phase of *F. graminearum* infection and colonisation. Given the increase susceptibility to FHB in response to JA and ethylene, this supports the JA/ ethylene pathway antagonism to the biotrophic phase of FHB. However, since the opposite effect of JA and ethylene was observed on FRR, we speculate that *F. graminearum* is adopting a more necrotrophic lifestyle on roots. This difference in trophism between tissues is not well documented. Differences in trophic state have been observed for other plant pathogens. Marcel and colleagues (2010) found that *M. oryzae* behaves as a biotroph during rice root infection despite adopting a hemibiotrophic lifestyle on leaves (Marcel et al., 2010). Hypothetical reasons for switching to necrotrophy may be due to a demand for a change in the resource acquisition method, the time required to overcome defences, or a change in plant defence response (Kabbage et al., 2015, Zeilinger et al., 2016). One or more of these factors may be causing this difference in fungal lifestyle between roots and shoots in the *B. distachyon* - *F. graminearum* pathosystem. Wang and colleagues (2015) found that wheat Sumai3 roots were susceptible to FRR despite this variety being known for its potent FHB resistance (Wang et al., 2015). Furthermore, Lyons et al. (2015) identified differing expression patterns between root and leaf infection by *Fusarium oxysporum*. These examples suggest that a unique defence mechanism might be occurring in roots which promotes a different trophic lifestyle in Fusarium.

**Fig. 5:**
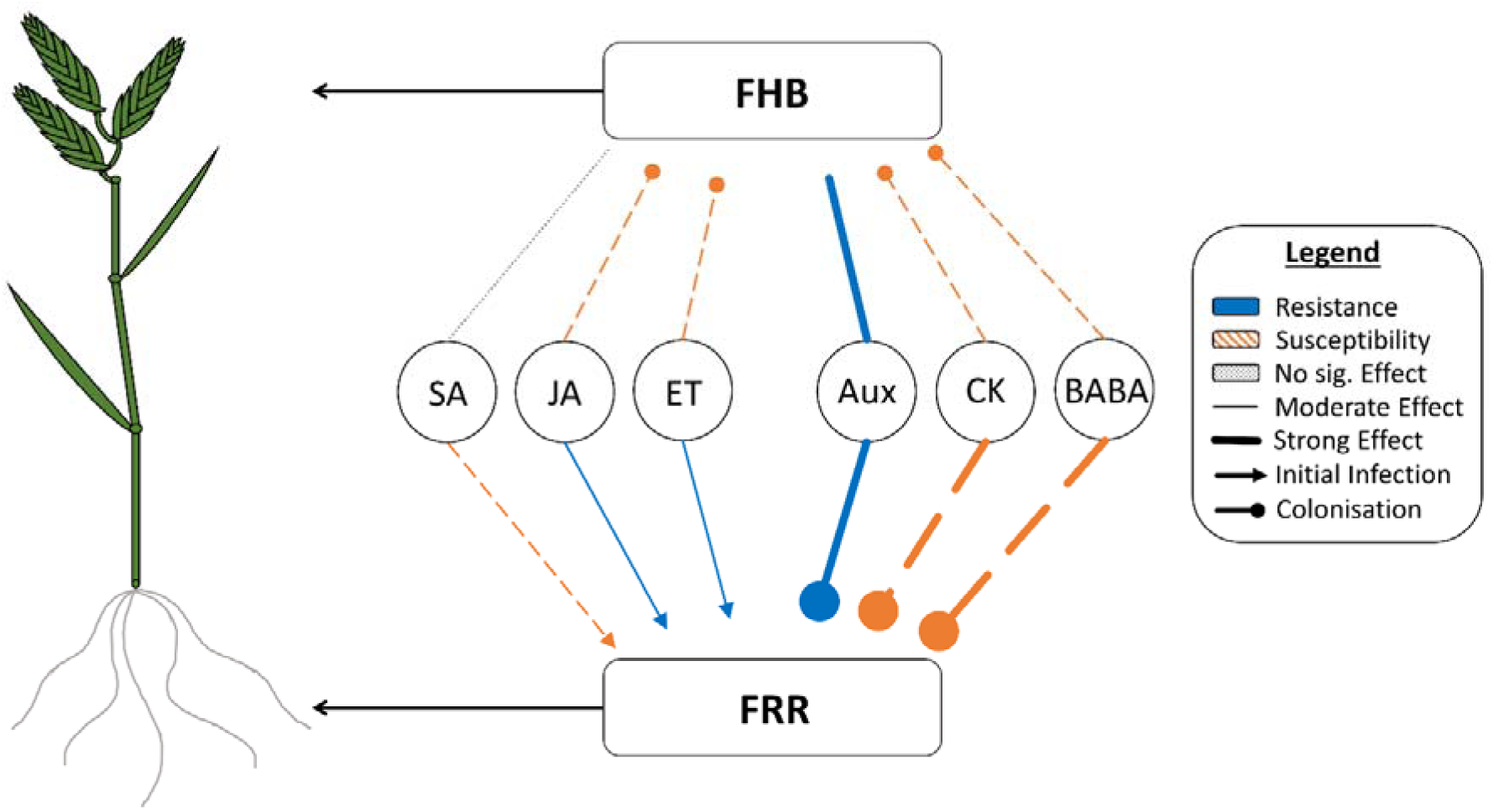
Model summarising all exogenous hormone application experiments (Fig. 1-5). Compares the effects between FHB and FRR in *B. distachyon* (on the left). This summary is a generalisation of the trend over time and includes different arrow thicknesses based on the potency of response caused by each phytohormone. Includes the predicted slope regression groups for each phytohormone presented as different arrow caps (excluding auxins and SA as these were not measured or not significantly different to control). Novel abbreviations: ET (ethylene), Aux (auxins), CK (cytokinins).

Contrary to the classic defence hormones, the development-related phytohormones induced similar effects on FHB and FRR (Fig. 5). Auxin application resulted in greatly increased resistance in both FHB and FRR in *B. distachyon*. Like the effect of auxins, the cytokinins trans-Zeatin and BABA also functioned in a tissue-independent manner, but they greatly increased susceptibility to FHB and FRR (Fig. 5). Aside from roles in plant defence, BABA can also affect plant development through reduction of growth and endogenous iron content (Wu et al., 2010, Koen et al., 2014). Therefore, BABA could be functioning similarly to the classic development-associated hormones auxin and cytokinin. There was a substantially larger effect of these three phytohormones on resistance compared to the classic defence phytohormones. We speculate that auxins, cytokinins, and BABA are affecting resistance in an SA/JA independent manner. In support of this, (Qi et al., 2016) found that exogenous application of IAA did not lead to an increase in endogenous levels of SA or JA. Furthermore auxin signalling independent of JA and SA was found to be important for *A. thaliana* resistance to two other necrotrophic pathogens (Llorente et al., 2008). In another example, elevated expression of a the IAA-amido synthetase gene *GH3-8* that regulates IAA levels promoted resistance to *X. oryzae* in rice independently of SA and JA (Ding et al., 2008). Cytokinins were also found to improve *N. tabacum* resistance to *P. syringae* in an SA and JA independent manner (Großkinsky et al., 2011). BABA functions synergistically with the abscisic acid defence pathway (Ton and Mauch-Mani, 2004) and there is evidence suggesting that abscisic acid promotes FHB susceptibility (Buhrow et al., 2016, Qi et al., 2016). The different effect of development hormones may be related to either phytohormone homeostasis, antimicrobial production linked to metabolic pathways of development hormones, or trade-offs with growth (Kazan and Manners, 2009, Huot et al., 2014, Albrecht and Argueso, 2017). Additionally, the opposite and marked response observed between cytokinins and auxins may be due to an antagonism between the two response networks (Naseem and Dandekar, 2012, Naseem et al., 2012). Overall, the effect of development-associated hormones on FHB and FRR may depend on their influence on basal resistance or specific growth/defence trade-offs independent of the trophic lifestyle. In contrast, the role of the classic defence hormones may be dependent on the trophic lifestyle of the pathogen.

Since multiple time points for both FHB and FRR were recorded, we observed differences in slope regression lines between phytohormones. We were able to separate the responses to compounds into two broad groups. The first ‘parallel’ group are those compounds that induce a significant difference in resistance at the first time point, and that the difference remained similar at later time points. The phytohormones SA, JA, and ACC (Fig. 1A-C) had this effect on FRR whereas no phytohormone displayed this on FHB. This trend implies that these compounds have an immediate effect on initial *F. graminearum* infection, after which the absence of deviation relative to the first time point suggests that the effect on FHB colonisation was much reduced or absent. The second ‘divergent’ group are those that have minor effects at the first time point, but where the differential in resistance increased at later time points. The growth and development-associated phytohormones exhibited this in FRR assays: BABA (Fig. 1D), both auxins (Fig. 2A and Fig. 2B), and both cytokinins (Fig. 2C and Fig. 2D). This trend implies that these compounds have a minimal effect on initial infection but affect the ability of the host to restrict *F. graminearum* colonisation. These trends are summarised in Fig. 5. Together the evidence supports the idea of fundamental differences in the functioning of growth/development associated phytohormones compared to the classic defence phytohormones in resistance to FRR. Secondly there were differences between FHB and FRR in response to the classic defence hormone. This supports our view that for FHB there is a period of biotrophic growth before switching to necrotrophy whereas in FRR this initial phase is much reduced or absent.

A model was produced to summarise the effects of the tested phytohormones on FHB and FRR in *B. distachyon* (Fig. 5). Phytohormones altered FRR and FHB resistance in *B. distachyon*. The results reflect the generally antagonistic nature of SA to JA/ ethylene in the two diseases. However they also show that these defence hormones induce disease-specific effects between FHB and FRR. Unlike the classical defence-related hormones, the development-associated hormones auxin and cytokinin had large tissue-independent effects on *B. distachyon* response to FHB and FRR, highlighting an important role of these hormones in the *B. distachyon* - *F. graminearum* pathosystem. Furthermore our data suggest that each hormone functions at different phases with some affecting on initial *F. graminearum* infection and others playing a greater role during colonisation. Future experiments are required to examine the host and pathogen transcriptome response during both FHB and FRR to provide additional evidence on the mode of trophism within infected tissues and gene expression of both partners in the interaction.

## Abbreviations

ACC: 1-aminocyclopropane-1-carboxylic acid;
BABA: 3-aminobutanoic acid;
dpi: days post inoculation;
FHB: Fusarium head blight;
FRR: Fusarium root rot;
IAA: indole-3-acetic acid;
JA: jasmonic acid;
NAA: 1-napthaleneacetic acid;
PDA: potato dextrose agar;
RNL: root necrosis length;
SA: salicylic acid.

## Acknowledgments

This work was supported by the BBSRC (grant number: BB/M011216/1) and BASF SE at Limbergerhof in Germany as part of the PhD studentship of J.F. Haidoulis. The authors would like to thank Dr Egon Haden and Dr Sebastian Rohrer from BASF SE for scientific advice, for providing materials including Bd21 and *Fusarium graminearum* Li600 isolate, and glasshouse space.

## Supplementary Figure Captions

<u>Fig. S1</u>: The visual decrease in RNL after IAA application (B) compared to control (A) at 6 dpi.

<u>Fig. S2</u>: The visual increase in RNL after Zeatin application (B) compared to DMSO control (A) at 6 dpi.

<u>Fig. S3</u>: The change in *F. graminearum*-induced FRR necrosis after application of gibberellic acid (A) and epibrassinolide (B) on Bd3-1 seedling roots.

<u>Fig. S4</u>: The visual effect of 10 µM abscisic acid on 7-day old Bd3-1 roots in the absence of inoculation.

<u>Fig. S5</u>: The change in number of *F. graminearum*-infected florets after four doses of salicylic acid 24 h before, 30 min before inoculation, 3 dpi, and 7 dpi on Bd3-1.

<u>Fig. S6</u>: The growth (measured as radius from mycelial epicentre) of *F. graminearum* on hormone-amended PDA at 2 dpa.

<u>Fig. S.7</u>: *F. graminearum* PH1 mycelial growth on PDA after 3 dpa with 8 mM IAA and NAA treated filter disks.

