## Supplementary Material for "Different effects of phytohormones on Fusarium head blight and Fusarium root rot resistance in *Brachypodium distachyon*"

These seven figures supplement the following manuscript:

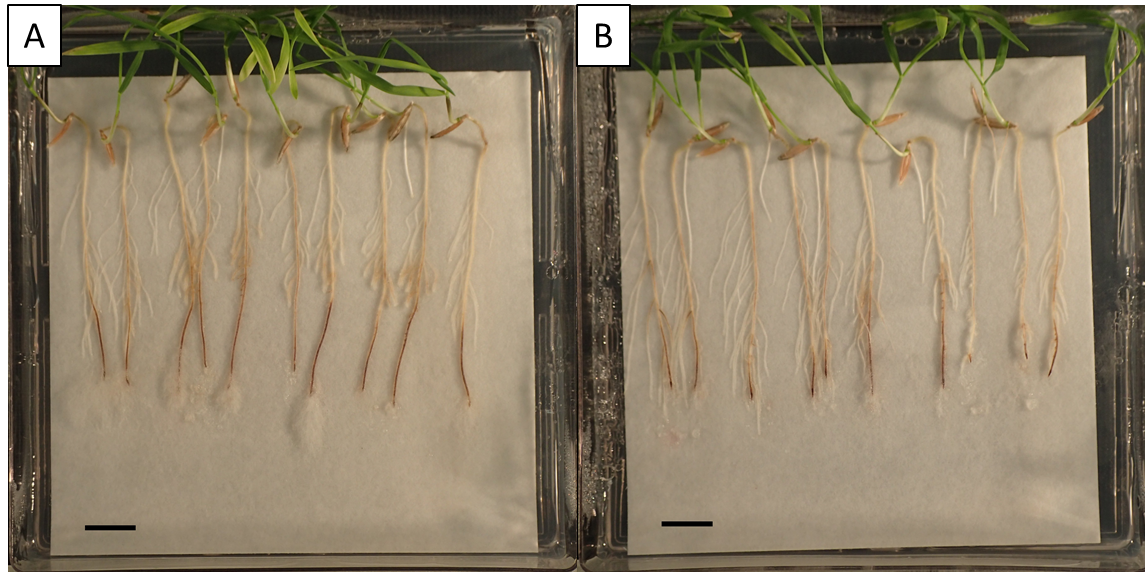

Supplementary Figure S1: The decrease in *F. graminearum*-induced root necrosis on Bd3-1 after IAA (Indole-3-acetic acid) application (**B**) compared to the control (**A**) at 6 dpi. Scale bars = 1 cm.

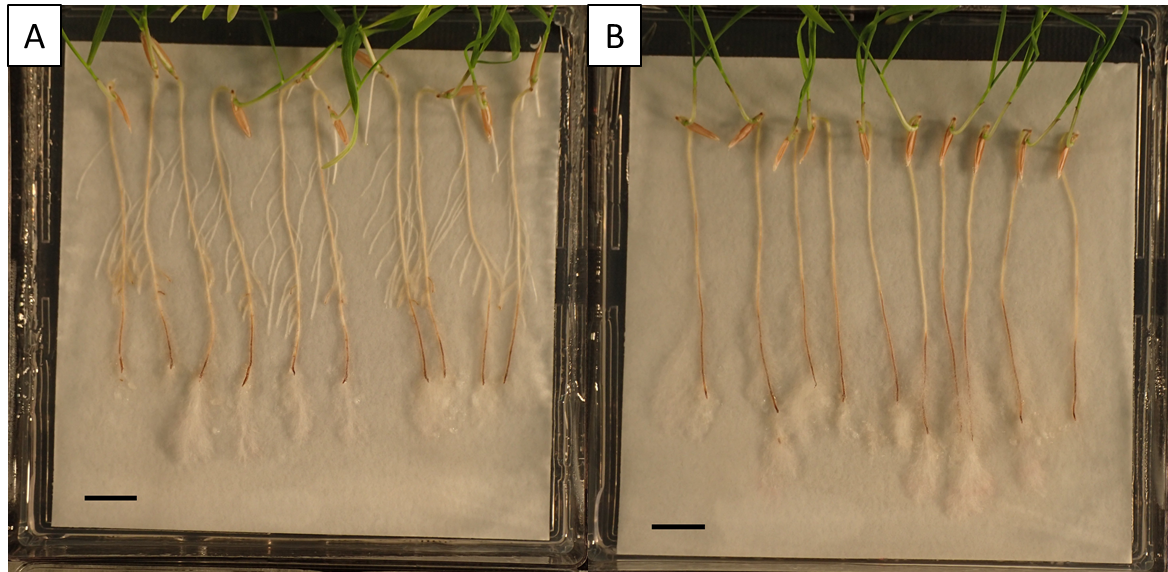

Supplementary Figure S2: The increase in F. graminearum-induced root necrosis on Bd3-1 after trans-Zeatin application (**B**) compared to the DMSO control (**A**) at 6 dpi. Scale bars = 1 cm.

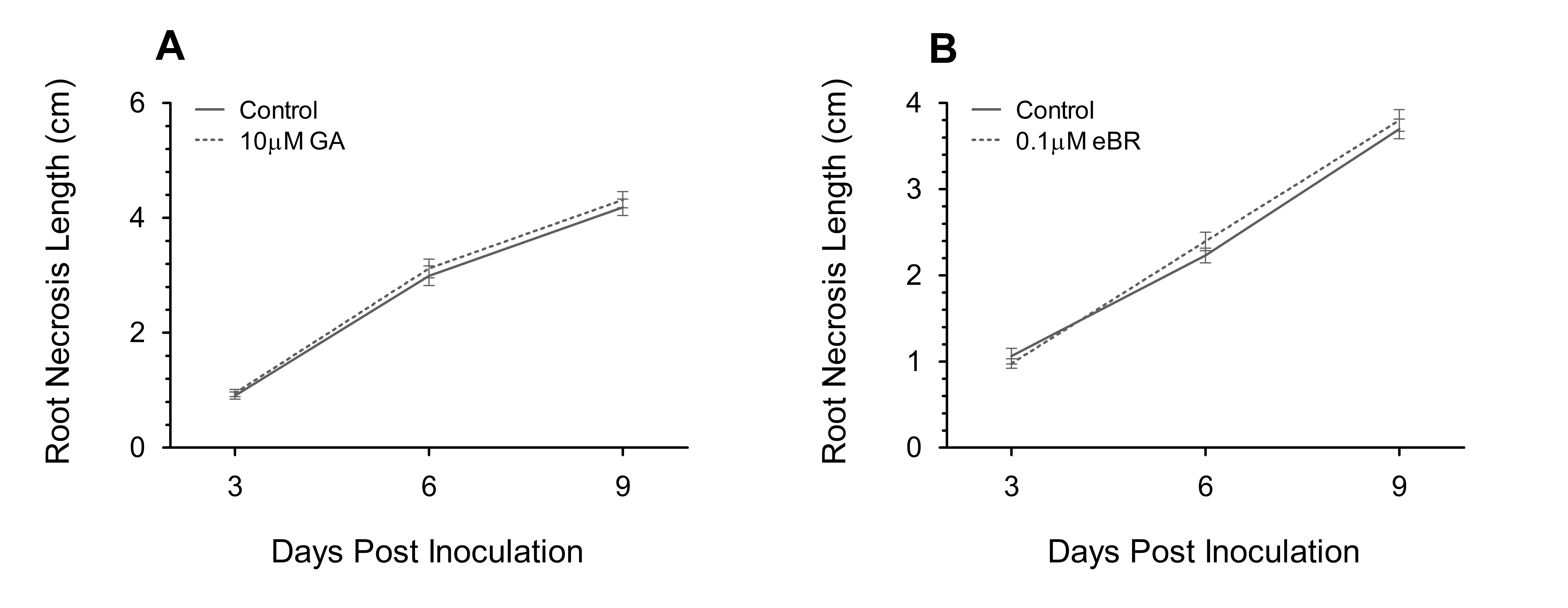

Supplementary Figure S3: The change in *F. graminearum*-induced root necrosis change on Bd3-1 seedling roots after application of gibberellic acid (GA) (**A**) and epibrassinolide (eBR) (**B**). (**A**) Control and GA treatments are both a water-based solvent treatment whereas (**B**) control and eBR are both a water-based treatment with 0.1 % ethanol solvent. Each data point is the mean root necrosis length ± SE (N = 28 (**A**), N = 30 (**B**) per treatment) each from one independent experiment. Generalised linear model-ANOVA significance levels comparing treatments were (**A**) p = 0.641 (3 dpi), p = 0.609 (7 dpi), and p = 0.53 (11 dpi), and (**B**) p = 0.579 sqrt transformed (3 dpi), p = 0.199 sqrt transformed (7 dpi), and p = 0.539 (11 dpi).

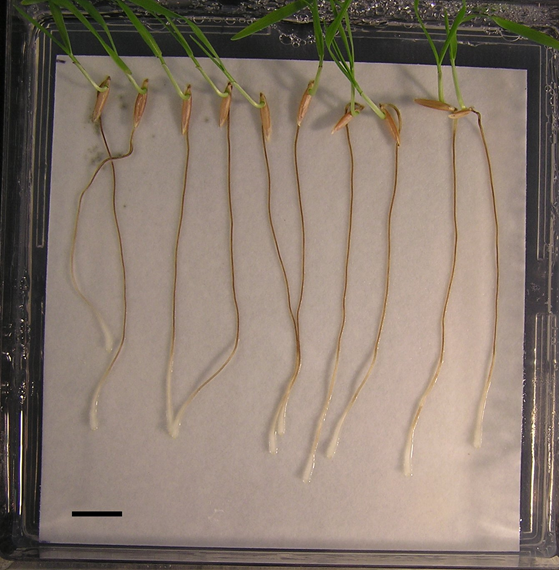

Supplementary Figure S4: The effect of 10 µM abscisic acid on 7-day old Bd3-1 roots in the absence of inoculation. Scale bar = 1 cm.

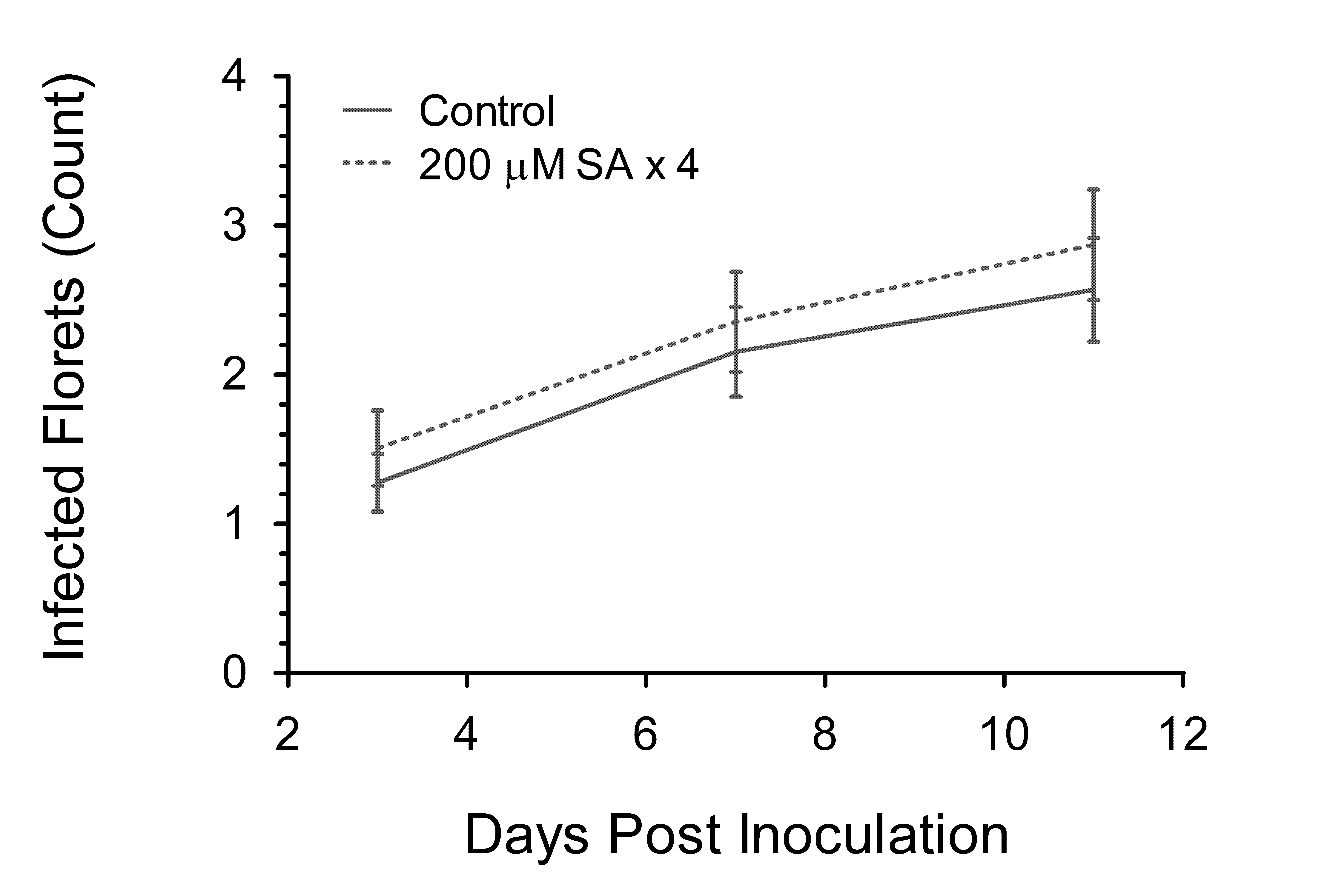

Supplementary Figure S5: The change in number of *F. graminearum*-infected Bd3-1 florets after four doses of salicylic acid (SA) 24 h before, 30 min before inoculation, 3 dpi, and 7 dpi. Each data point is the mean number of florets infected ± SE (N = 65 control, N = 63 SA) from one independent experiment. Generalised linear model-ANOVA significance levels comparing treatments were p = 0.457 (3 dpi), p = 0.979 sqrt transformed (7 dpi), and p = 0.616 sqrt transformed (11 dpi).

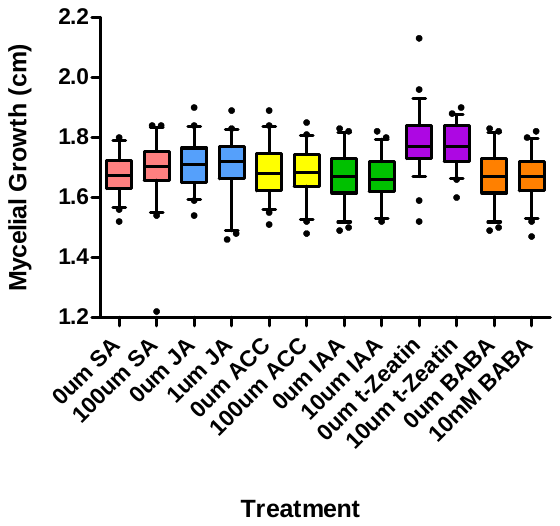

Supplementary Figure S6: The growth (measured as radius from mycelial epicentre) of F. graminearum PH1 on hormone-amended PDA at 2 dpa (days post application). Each data point is the average of approximately 46 measurements from 12 biological replicates (4 measurements per biological replicate) ± SE. A 95^th^ percentile whisker end cap was used for all treatments. Generalised linear model-ANOVA comparing compound to respective control; p = 0.247 (SA), p = 0.364 (JA), p = 0.918 (ACC), p = 0.568 (IAA), p = 0.729 (BABA), p = 0.475 (tZ). Abbreviations; SA (Salicylic acid), JA (Jasmonic acid), ACC (1-aminocyclopropane-1-carboxylic acid), and BABA (3-aminobutanoic acid), tZeatin (trans-zeatin).

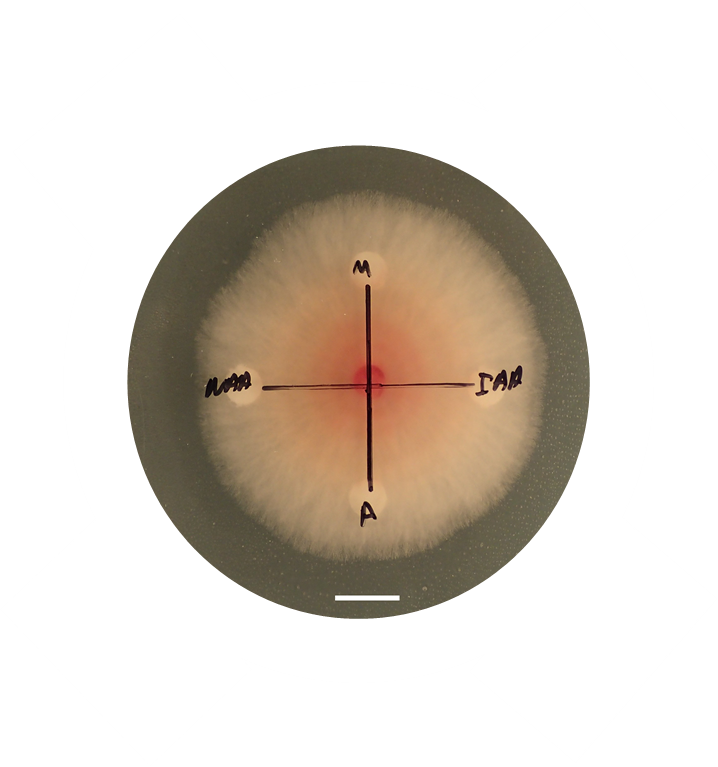

Supplementary Figure S7: F. graminearum PH1 mycelial growth on PDA with 8 mM Indole-3-acetic acid (IAA) and 1-Naphthaleneacetic acid (NAA) treated filter disks after 3 dpa. Image is one of three Petri-dishes. The experiment was repeated twice. IAA was dissolved in Methanol (M) whereas NAA was dissolved in Acetone (A). Scale bar = 1 cm.

All scale bars for images were made with ImageJ: (Abràmoff *et al.*, 2004)

**Abràmoff MD, Magalhães PJ, Ram SJ**. 2004. Image processing with ImageJ. Biophotonics international **11**, 36-42.
